# Passive droplet microfluidic platform for high-throughput screening of microbial proteolytic activity

**DOI:** 10.1101/2024.06.04.597455

**Authors:** Luca Potenza, Łukasz Kozon, Lukasz Drewniak, Tomasz S. Kaminski

## Abstract

Traditional bacterial isolation methods are often costly, have limited throughput, and may not accurately reflect the true microbial community composition. Consequently, identifying rare or slow-growing taxa becomes challenging. Over the last decade, a new approach has been proposed to replace traditional flasks or multi-well plates with ultrahigh-throughput droplet microfluidic screening assays. In this study, we present a novel passive droplet-based method designed for isolating proteolytic microorganisms, which are crucial in various biotechnology industries. Following the encapsulation of single cells in gelatin microgel compartments and their subsequent clonal cultivation, microcultures are passively sorted at high throughput based on the deformability of droplets. Our novel chip design offers a 50-fold improvement in throughput compared to previously developed deformability-based droplet sorter. This method expands an array of droplet-based microbial enrichment assays and significantly reduces the time and resources required to isolate proteolytic bacteria strains.

## INTRODUCTION

Proteolytic microorganisms play a crucial role in modern biotechnology due to their ability to break down proteins. Proteolytic activity holds a wide range of applications in industries such as bioenergy production, food processing, detergent production, and pharmaceuticals^1–4^. Proteolytic bacteria also play a role in waste management and biomedical diagnostics. Proteases produced by these strains are widely used as biocatalysts because they facilitate environmentally friendly chemical reactions under mild conditions^5,6^. In microbiology, the characterization of microbial proteases is based on the isolation of proteolytic microorganisms on selective media^7^.

One of the most common techniques to detect microbial proteolytic activity is cultivation on agar plates containing skimmed milk. Together with lactose, the amino acids from dairy proteins are carbon sources in this agar medium. The proteolytic activity is indicated by a transparent halo around colonies on the milk-white background that appears exclusively around the proteolytic colonies^8^. Nonetheless, this assay can lead to false positive results, as clear halos could be attributed to lactose utilization and the subsequent production of acid metabolites rather than being caused by the proteases produced by a proteolytic strain^9^. Moreover, this protocol presents limitations such as low throughput and the necessity for cultivation on a solid medium. Several studies have shown how growing bacteria on agar can negatively affect the recovery of certain microorganisms, thus reducing the overall screening capabilities^10^.

Numerous techniques are employed to quantify protease activity, involving various substrates, both natural and synthetic, and coupled with colorimetric readouts. Common colorimetric assays involve using protease substrates such as azocasein^11^ or chromogenic tripeptides^12^ that are catalyzed to products that can be detected via absorbance. In the alternative method, the ninhydrin-based method exploits the reaction between ninhydrin and amino acids released during protein breakdown. In this assay, unreacted ninhydrin turns yellow, while amino acid-ninhydrin complexes turn purple. The intensity of the purple color allows for the determination of amino acid concentration resulting from protein degradation^13^. Despite being widely used, classical protocols for screening microbial proteolytic activity have several limitations. Their sensitivity is too low for accurate detection in cases where proteolytic microbes are present in low quantities, have slow growth rates, or produce proteases with low activity. Even with a transition to a low-volume multi-well plate format, many assays can be costly and time-consuming, which becomes a bottleneck with a large number of samples.

Microfluidic assays offer a distinct advantage by enabling the encapsulation, incubation, analysis, and sorting of individual microbial cultures within controlled microenvironments^14^. Various studies demonstrated the utility of droplet microfluidics for screening and enriching microbes, for example, based on their growth,^15^ production of antibiotics^16^ or biocatalytic activity^17^. Detection of proteolytic activity in a droplet-based format has been described only for single mammalian cells, and all these methods are based on fluorescence readouts^18–20^.

In contrast to the above-mentioned protocols, passive methods for microfluidic enrichment of microorganisms leverage the advantages of passive microfluidic sorting. This approach eliminates the necessity for complex instrumentation, thereby simplifying designs, reducing operational expenses, and enhancing user-friendliness. Utilizing phenomena such as surface tension^21^, visco-elasticity^22^ and difference in size or density^23^, passive systems enable selective sorting of droplets. A recent study introduced a system employing passive screening to enrich agarolytic bacteria. The method involved encapsulating microbial cells in microdroplets containing a hydrogel. As the hydrogel gradually degraded, the droplets became more deformable. A double rail-driven microfluidic design was utilized to passively sort microdroplets. Despite the successful sorting of droplets containing agarolytic microorganisms, the method exhibited a limited throughput, not exceeding a single droplet per second^24^, which is too low for the enrichment of rare strains from complex microbial consortia.

In this study, we thoroughly explored the utilization of a droplet-based system within the field of microbiology, with a specific focus on the selection of proteolytic microbial strains. Key aspects of our research include: i) the design of a passive microfluidic droplet sorter, ii) the utilization of gelatine droplets for the clonal cultivation of microbes, and iii) the establishment of a novel protocol for single-cell studies and microbial isolation. Validation of the efficacy of the deformability-based passive droplet sorter (DPDS) was conducted through an enrichment test to isolate proteolytic strains with exceedingly low abundance, within a mock microbial community. Our efforts led to the development of a passive microfluidic workflow demonstrating competitive accuracy and, to some extent, throughput compared to active systems, which typically employ more complex screening methods such as fluorescence-^16^ or absorbance-based^25^ assays.

## MATERIALS AND METHODS

### Bacteria cultivation

During the development and validation of our microfluidic method, we utilized two different strains: i) The *Pseudomonas aeruginosa* strain was used to generate positive droplets containing a high proteolytic bacterium, and ii) the non-proteolytic *Escherichia coli DH5α* strain served as a negative control. To prepare overnight cultures, we inoculated a single colony of each strain into 10 ml of LB liquid medium in flasks, which were then incubated at 30°C overnight while shaking at 100 rpm. To assess the microbial growth, we measured the optical density at 600 nm (OD_600_) of 200 μl of each cell culture using a plate reader (Sunrise, Tecan Trading AG) and MagellanTM software. The concentration of cells and the calculations to achieve the desired cell/droplet occupancy were based on OD_600_ readings from the overnight LB culture. The droplet cultivation medium comprised two stock solutions to be diluted: a gelatin stock (300 g/L gelatin) and an LB medium (LB 1x). The gelatin stock was preheated to 40°C to liquefy it. Then, the gelatin was diluted with a 0.9% NaCl saline solution to achieve a final concentration of 75 g/L (7.5% gelatin). The LB needed to be mixed thoroughly with the gelatin solution to obtain the final medium composition: LB 0.5x supplemented with 7.5% gelatin. To prevent foam formation, bacteria were first mixed with LB by vortexing and then gently combined with the gelatin solution through careful pipetting up and down.

### Fabrication of microfluidic devices

Masks with microfluidic chips were designed in AutoCAD 2021 (Autodesk) and then printed onto a high-resolution film mask (Micro Lithography Services Ltd). The CAD files containing the designs of a droplet sorting device can be found as a supporting .dxf file. A detailed protocol for microfabrication of devices is described in the Supporting Information (SI). Briefly, master molds were made by photolithography via the coating of silicon wafers with SU-8 2025 photoresist (Kayaku Advanced Materials) and exposure to UV light using film masks and an MJB-4 mask aligner (SUSS MicroTech). The same mask aligner was used for aligning the microfilm mask and UV exposure of the second layer. Replicas of PDMS devices were next fabricated by soft lithography by casting polydimethylsiloxane (PDMS, Sylgard 184, Dow Corning) on master molds. Once PDMS solidified, the inlet and outlet holes were punched in the chips, and then the devices were bonded to glass slides via an automated plasma chamber (Zepto, Diener). A hydrophobic coating of microfluidic channels was performed by flushing the chips with 1% trichloro(1H,1H,2H,2H-perfluorooctyl)silane (Sigma-Aldrich) in Novec HFE-7500 (3M) and placing them on a hot plate (80°C) for 1 hour.

### Gelatin droplet generation and cultivation of encapsulated microbes

Bacterial cells were resuspended from an LB overnight culture into the rich-gelatin medium to achieve the desired cell concentration. The resulting bacterial suspension was next aspirated into a 1 mL glass syringe (gastight series, Hamilton). Microemulsions were generated in a flow-focusing droplet generator (50 x 50 μm) with flow rates of 1.08 mL/h for the oil phase composed of Novec HFE-7500 (3M) with 2.5% 008-FluoroSurfactant (RAN Biotechnologies) and 0.6 mL/h for the cell suspension in LB-gelatin medium at a throughput of 1.5-2 kHz. The room temperature for droplet generation was set at 25°C. Approximately 150 μL of droplet emulsion was collected inside a droplet chamber^26^ and incubated at 40°C for 72 hours. A peristaltic pump was employed (Reglo ICC, Ismatec) to provide oxygen dissolved in the oil to the bacteria encapsulated inside droplets, while preventing air bubbles from entering the droplet chamber. The flow rate of oil flowing through the emulsion during incubation was set to 0.6 mL/h. Detailed protocol for the oxygen supply system is described in the Supporting Information (SI).

### Passive droplet sorting

Three glass syringes (gastight series, Hamilton) were filled with HFE-7500 oil, attached to PTFE tubing, and locked into position on syringe pumps (Nemesys, Cetoni). A first 0.5 mL syringe was filled with 1% RAN fluorosurfactant in oil and used as the first spacing oil to prevent droplet merging in the re-injection chamber. A second 1 mL syringe was filled up with pure oil and was employed as the second spacing oil. Similarly, a third 10 mL syringe was filled with pure HFE-7500 oil that was used as a sorting oil. The top outlet tubing of the droplet incubation chamber was inserted into a 1 mm inlet to reinject droplets in the sorter device. The tubing for spacing and sorting oils were also connected to their respective inlets. The syringe pumps simultaneously pushed oils and droplets through the device. The flow rates required for sorting proteolytic microcultures within droplets were as follows: 3·10^−2^ mL/h for the first spacing oil, 6·10^−2^ mL/h for the second spacing oil, and 3 mL/h for the sorting oil. The emulsion flow rate was set at 1.5-1.8·10^−2^ mL/h. Videos were captured using a high-speed camera (AX-100 Mini, Photron) and an inverted optical microscope (IX73, Olympus).

### Determination of enrichment factor

Proteolytic strain microcolonies confined in droplets were collected in sterile 1.5 mL Eppendorf tubes. Both the collection tube and the tubing connecting the tube with the microfluidic chip were autoclaved at 121°C for 15 minutes before the experiment. Passive sorting of proteolytic microbes was carried out for 30 minutes, after which the tubing was disconnected from the positive outlet of the sorter and flushed with filtered and sterile HFE-7500 oil. The volume of oil in each sample was adjusted to 250 μL, and the emulsion was disrupted by adding 62.5 μL of 1H,1H,2H,2H-perfluoro-1-octanol (PFO) (Alfa Aesar) and 200 μL of sterile 0.9% NaCl saline solution. The two-phase mixture was first vortexed for 90 seconds and then centrifuged for 60 seconds at 2000 rpm to better separate the different phases. The aqueous phase was used to plate the enriched bacteria on Petri dishes containing agar supplemented with 30g/L of skimmed milk powder. The plates were incubated at 40°C for 24 hours, and then examined by visually counting the number of colonies. Colony-forming units were counted for each plate and dilution, distinguishing between the negative colonies (no transparent halo around the colonies) and the positive ones (exhibiting a transparent halo).

### Data analysis

To assess the performance of droplet sorting in our microfluidic module, slow-motion videos were examined using ImageJ software to calculate the displacement factor values. Initially, an average background image was created from the video by using the ‘Image/Stack/Z Project’ tool in ImageJ, using the Average Intensity projection type. Next, this background image was subtracted from each frame of the video using the ‘Process/Image Calculator’ function in ImageJ. By doing so, we aimed to isolate the droplets from the rest of the image, which would otherwise include channels and chip structure. Therefore, to identify the droplets in the resulting ‘background-subtracted’ video, the ‘Image/Adjust/Threshold’ function in ImageJ was utilized. This function set a threshold to distinguish the droplets’ boundaries based on their pixel values. The binarized edges of the droplets were then subjected to the ‘Fill holes’ function (‘Process/Binary/Fill Holes’) so that the resulting masks would also include the interior of the droplets. The coordinates of the centres of the sorted droplets were then extracted from each frame of the video slice. This operation was done by utilizing the ‘Set Measurements’ and ‘Analyze Particles’ functions with the ‘Centroid’ and ‘Overlay’ options selected and the ‘size (pixel^2)’ filter was set to enumerate exclusively moving particles with an area above 8500. The raw data was further analyzed and visualized in RStudio and Python.

## RESULTS AND DISCUSSION

A well-established method for screening microbial proteases in bulk is based on the use of a gelatin-rich medium^27^. Gelatin, a product of the partial hydrolysis of collagen in animal tissues, serves as both a substrate for microbial proteolytic activity and a solidifying agent. Proteases enzymatically break down the gelatin matrix, increasing solubility by converting proteins into smaller, soluble components. This process disrupts the gelatin structure, reducing its gel-like properties. As a result, when a proteolytic strain is cultivated in gelatin, the medium liquefies. Utilizing the significant change in the deformability properties of gelatin droplets, we propose here a microfluidic method for selecting proteolytic microbes comprising four consecutive steps, as illustrated in **Figure 1**. First, single cells are encapsulated inside picoliter droplets via a flow-focusing device. Droplets are then collected and incubated off-chip inside droplet chambers to allow microbial growth. A constant flow of oil is provided by a peristaltic pump to ensure the oxygenation of droplets. To prevent the accumulation of air bubbles in the chambers, a bubble trap tube is positioned between the pump and the droplet chamber. Droplets are next reinjected into the deformability-based passive droplet sorter (DPDS) device, where solid droplets (negatives) are pushed along the barrier, while more liquid droplets (positives) are squeezed underneath it and collected into the positive outlet. The enriched microbial cultures encapsulated in droplets are subsequently cultured on a solid selective medium to confirm their proteolytic activity and the enrichment factor of the protocol.

**Figure 1.**
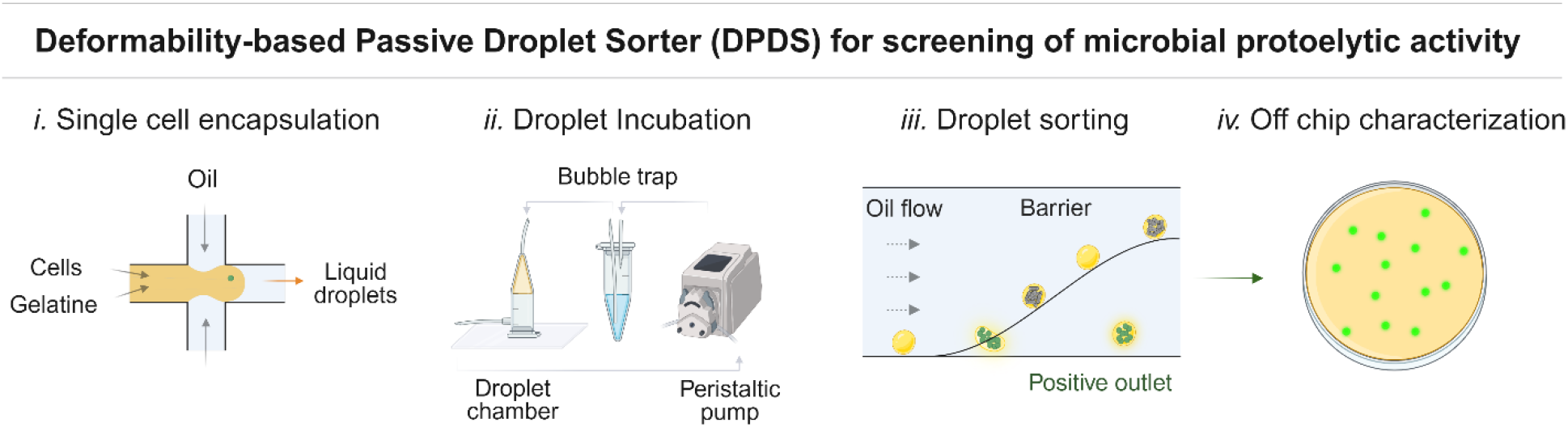
Schematic of the high-throughput passive screening of proteolytic microbial activity. The microfluidic assay is divided into four consecutive steps. i) single bacterial cell encapsulation into liquid gelatin droplets, ii) droplet incubation in dynamic conditions, iii) sorting of droplets based on their deformability properties, ii) off-chip characterization of sorted colonies using standard microbiology methods.

The development of the method for selecting proteolytic bacteria involved multiple stages: i) designing and validating a passive sorter which selects droplets based on deformability, ii) defining the parameters for efficient droplet sorting, iii) optimizing bacterial growth conditions in picoliter droplets, and iv) demonstrating the enrichment of proteolytic bacteria from a mock microbial consortium.

### The design of the deformability-based passive droplet sorter

The layout of the double-layered passive sorter, depicted in **Figure 2**, represents a novel design which comprises several functional parts. Droplets flowing from the external incubation container first enter the re-injection chamber. The narrower part of the chamber where the 2nd layer of the chip transitions to a part with a height of 30-μm-high forces the droplets to be delivered as a monolayer for efficient spacing with oil. The chamber narrows to a 50-μm-wide main channel and intersects with a re-injection junction composed of a double flow-focusing delivering two consecutive streams of spacing oil. The scope of such a double-spacing approach is to gradually accelerate droplets, enhancing the overall throughput and simultaneously preventing droplet break-up and/or merging, a phenomenon which we observed for high flow rates of spacing oil and low concentration of gelatin in droplets. Once droplets are evenly spaced, they are pushed against a barrier by the sorting oil flowing from the branched tree layout. The barrier is 12.5-μm-thick and 30-μm-high element inclined at 35° to the flow direction. Proteolytic bacteria cultures degrade the gelatin protein matrix and this results in a change of deformability properties that allow positive droplets to squeeze underneath the barrier and be collected in the positive channel. On the other hand, empty droplets and those with non-proteolytic cultures remain solid, and they are pushed along the barrier to the negative channel and discarded.

**Figure 2.**
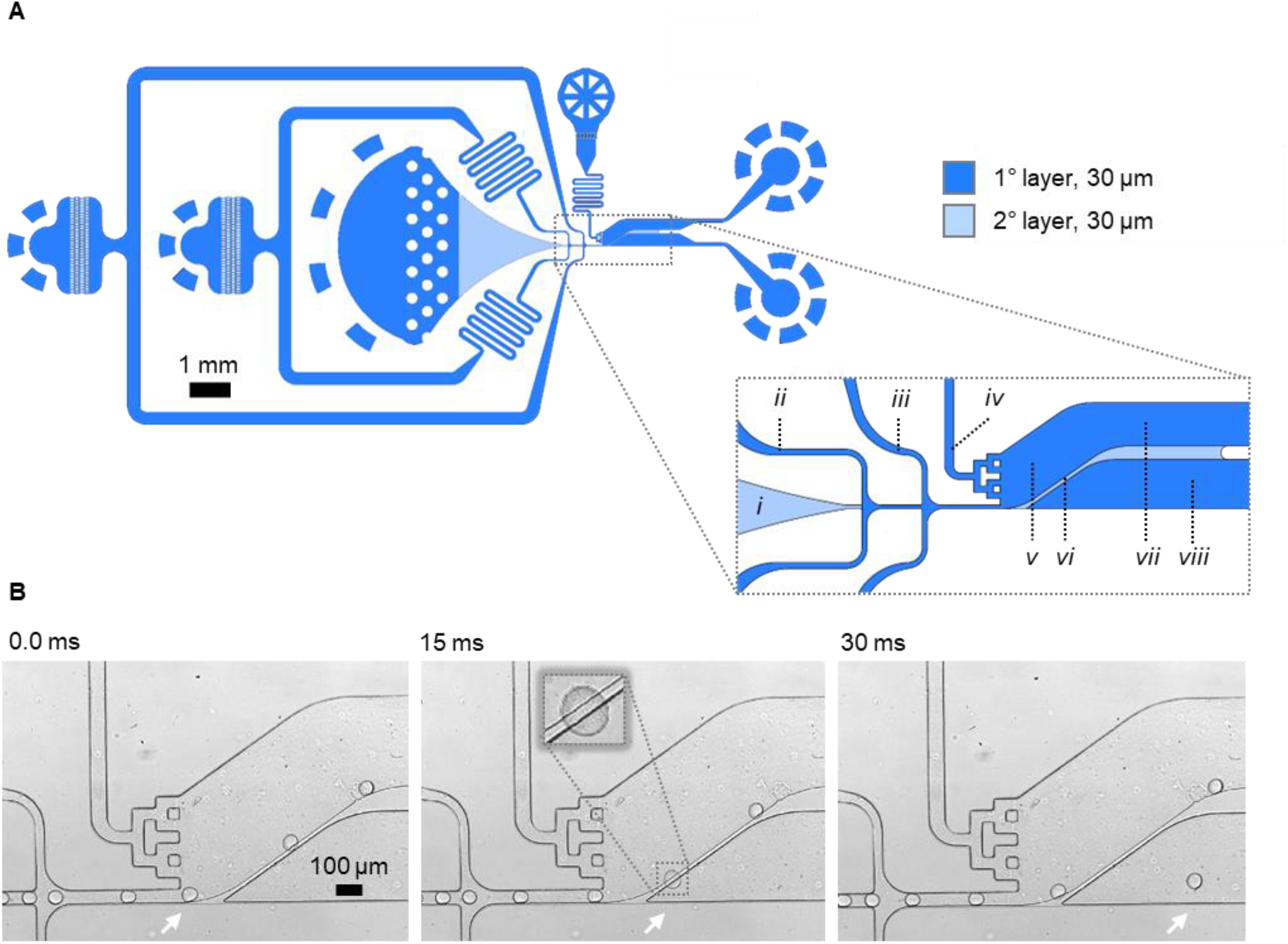
The design of the double-layered DPDS device. Panel A shows the architecture of the passive sorter, which includes: i) the droplets re-injection chamber, ii-iii) two channels for consecutive streams of spacing oils, iv) the sorting oil channel, v) the main chamber of the chip where sorting is executed, vi) the barrier that derails droplets, vii) the outlet for negative droplets located on the top part of the chamber, and viii) the outlet for positive droplets in the lower part of the chamber. Microphotographs showing the sorting event of a positive droplet are depicted in Panel B.

### Setting up conditions for generation and incubation of gelatin droplets

The assay we propose requires solid droplets to be effectively derailed by the barrier into the negative channel. As shown in the **Supporting video S1**, an aqueous solution of 7.5% gelatin is first partitioned into monodisperse emulsion generation (∼1.5-2 kHz, 100 pL) at 25°C using a flow-focusing junction. Next, these droplets are incubated at 40°C and subsequently sorted at 20°C. The room temperature during generation and screening should be controlled by the use of an air conditioning system or the implementation of a microscope environmental chamber. Despite achieving monodisperse droplet generation for a relatively high concentration of gelatin, we encountered droplet merging within 24 hours of incubation, which hindered the screening process To address this challenge, we investigated the relationship between static and dynamic droplet incubation^28^. We monitored droplets by transferring them from the incubation chamber to a counting chamber to track emulsion stability. Our findings revealed that dynamic droplet incubation (0.6 mL/h, with 5% RAN FluoroSurfactant in HFE-7500) effectively mitigated droplet instability for up to 96 hours of incubation. An additional observation from dynamic incubation was the gradual shrinkage of droplets over time – a phenomenon observed before for microbial droplet culture under oil flow^28^. We determined that droplets volume is reduced by approximately 20 pL after 96 hours under cultivation conditions. We accounted for the shrinkage phenomenon by intentionally generating droplets that were initially 100 pL in volume, aiming for an approximate volume of 80 pL on the day of screening.

### Influence of volume, gelatin concentration and temperature on the successful sorting of proteolytic cultures

The development of a barrier-driven sorting mechanism relies on the deformability of positive droplets that can be squeezed and collected in the positive outlet channel. Through our investigations, we have determined how three key parameters such as volume, gelatin concentration and temperature, significantly impact the sorting process. First, the volume of droplets is crucial in determining their sorting process. In our study, we assessed the relationship between droplet diameter and their impact on barrier-driven sorting within a single DPDS device, while maintaining a room temperature of 20°C. Various volume droplets composed of LB 0.5x medium without gelatin were subjected to deformability-based screening, as illustrated in **Figure 3**. We found that the optimal volume of droplets for achieving the best performance was 80 pL. Droplet volumes exceeding 100 pL resulted in a high rate of false negatives, as these droplets despite being positives, were too large to deform against the barrier and exit the chip via the positive channel.

**Figure 3:**
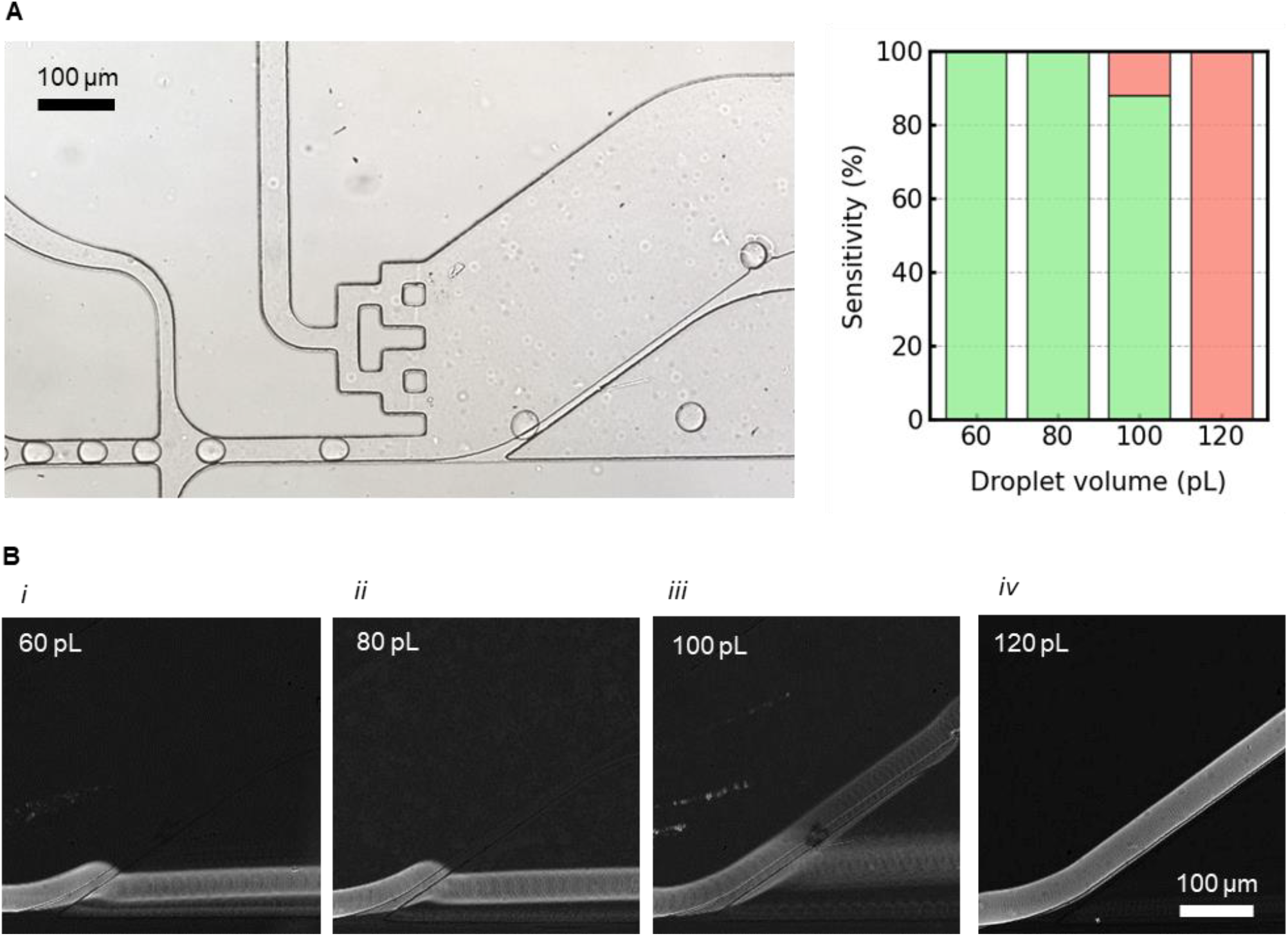
Influence of droplet volume on passive sorting. Panel A presents the impact of droplet volume on the displacement of positive droplets using four different types of LB 0.5x droplets. The column bars plotted demonstrate that the DPDS sorter is highly sensitive to droplet volumes below 100 pL. Green bars represent correctly sorted positive droplets, while red indicates positive droplets directed to the negative channel (false negatives). Panel B showcases sorting accuracy for different droplet volumes tested. Stack images, based on standard deviations, were generated from videos captured during the passive sorting process. During testing, the sorting oil flow rate ranged from 3-4.2 mL/h, and spacing oils were set to 3 ·10^−2^ mL/h and 6 ·10^−2^ mL/h. The room temperature was maintained at 20°C throughout the experiment.

Moreover, since gelatin droplets undergo solidification based on temperature, we had to evaluate the influence of gelatin concentrations on the sorting efficacy. We achieved accurate results under the slightly optimized temperature conditions, with a switch from 25ºC for droplet generation to 20ºC for 30-minute incubation prior to and during the execution of sorting. We examined six different gelatin concentrations (0, 15, 30, 45, 60, 75 g/L) as shown in **Figure 4**, monitoring the gel/liquid properties of the medium using tube samples placed beside the microfluidic rig. This straightforward experiment allowed us to quickly assess the feasibility of microfluidic screening at a room temperature of 20°C. The DPDS device is able to exclusively enrich droplets with 0% gelatin, while discarding all the droplets containing from 1.5% up to 7.5% gelatin concentration under optimum flow conditions for maximized throughput.

**Figure 4.**
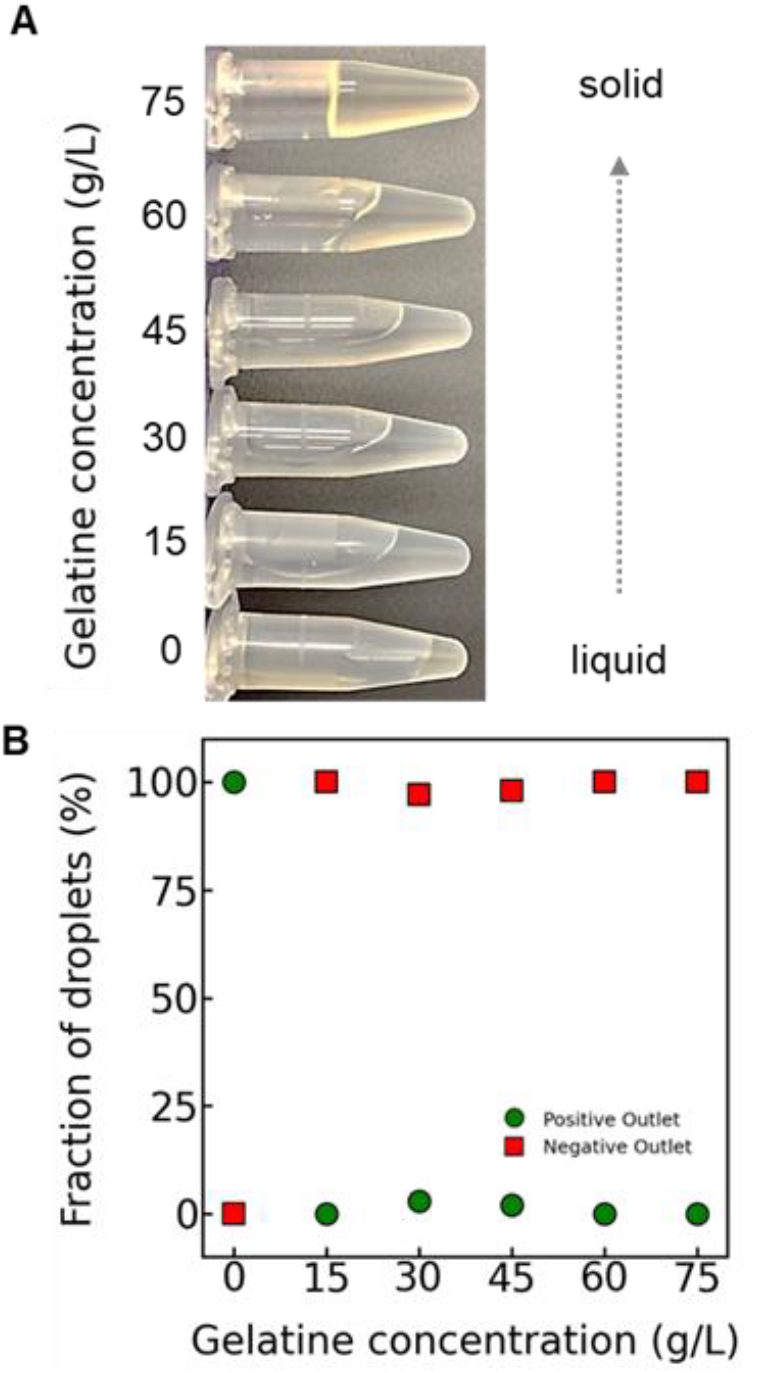
Influence of gelatin concentration and temperature on cultivation medium liquidity. Panel A: The image of various gelatin concentrations captured 30 minutes after introducing the tubes into the room where the experiment was conducted. The room temperature was maintained constant throughout the experiments using an air conditioning system. The data presented in Panel B, resulting from video analysis, shows a high level of accuracy in sorting exclusively the LB 0.5x droplets (no gelatin), whereas the barrier design under the above-mentioned conditions proves to be very effective in derailing droplets even with a minimum amount of gelatin.

### Microbial growth optimization - influence of cultivation medium and gelatin concentration

To ensure significant growth of both proteolytic and non-proteolytic strains, we tested various cultivation media, as reported in the Supporting Information (SI) section. Our data revealed that 2 times diluted LB medium (0.5x LB) supplemented by gelatin facilitated faster growth of bacteria both in droplets and in bulk. Although gelatin serves as a substrate for proteolytic microbes, high concentrations of gelatin tend to inhibit the growth of even the proteolytic *P. aeruginosa* strain, as shown in Supporting Information (SI) in **Figure S4**. Furthermore, concentrations exceeding 75 g/L of gelatin resulted in emulsion instability, leading to droplet merging during incubation. Consequently, we opted for a final medium composition of LB 0.5x supplemented with 75 g/L of gelatin (7.5% w/v), as illustrated in **Figure 5**. This composition consistently promoted the growth of the reference strains both in bulk and in droplet format. Additionally, droplets remain monodisperse during incubation, which facilitates quantitative screening of proteolytic activity.

**Figure 5.**
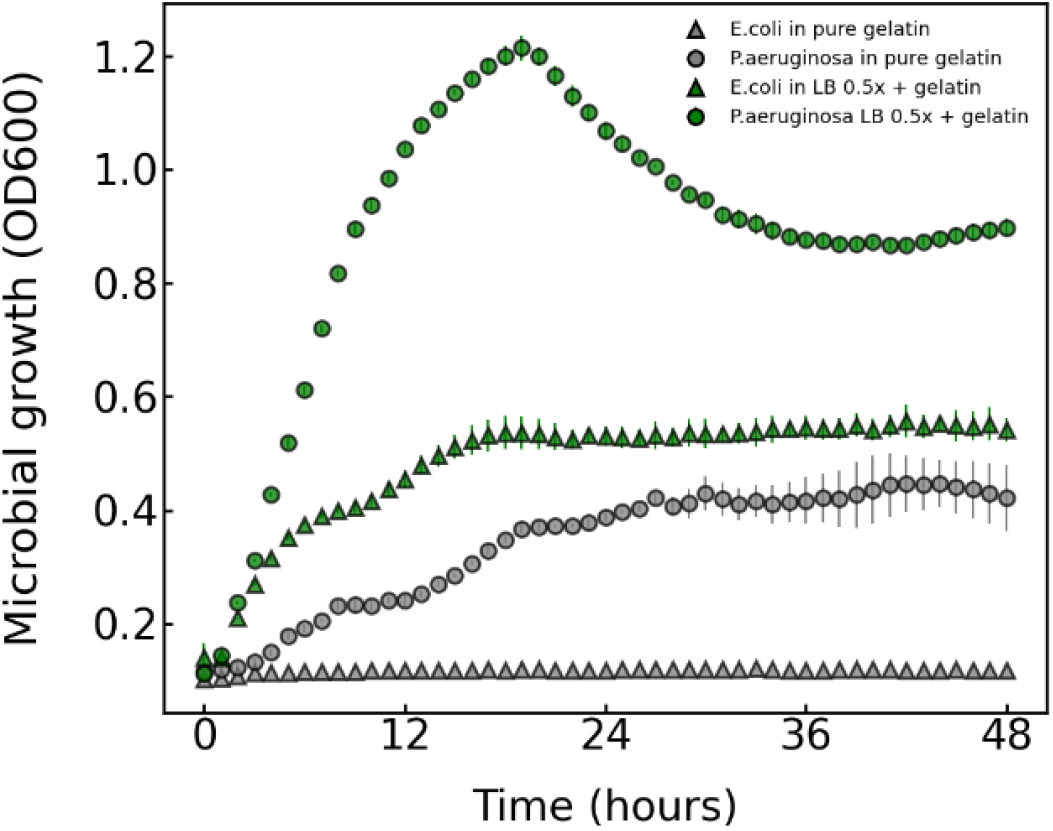
Cultivation medium optimization and growth of reference strains. The microbial growth of reference strains in pure gelatin 7.5% was compared against the medium chosen for in-droplet cultivation - LB 0.5x + 7.5% gel. The scatter plot illustrates the growth in bulk of both proteolytic and non-proteolytic bacteria utilized in the development of our microfluidic screening for proteolytic activity. Growth was monitored every hour using a well-plate reader set with shaking at 40°C.

### Optimizing flow parameters for efficient sorting

After several rounds of iterative design and testing, we optimized additional parameters that significantly impact the efficiency of the sorting process. The first tested factor was the droplet volume and the flow rates of both the emulsion and oils used in the sorter system. We used as a reference strain a proteolytic bacterium that was isolated from wastewater sludges and identified as *P. aeruginosa* as described in the SI section. Throughout the optimization process and the assessment of sorting accuracy, we employed two types of droplet populations. i) ‘Negative’ - empty and *E. coli* droplets represented a baseline reference. Given the non-proteolytic nature of the negative strain we selected, such droplets were found to be as negative as the empty ones despite consistent *E. coli* growth. ii) ‘Positive’ droplets contain *P. aeruginosa*. To generate microdroplets of uniform volumes, we employed a dedicated flow-focusing device with a 50×50 um junction. Microdroplets were collected within a simple droplet chamber, which facilitates easy re-injection and further manipulation^26^.

Next, we proceeded to test different flow rates of sorting oils to determine parameters to achieve optimal sorting accuracy and throughput. The purpose was to determine the numbers of both negative and positive droplets in true and false sorting events. As reported in **Figure 6**, we detected positive (*P. aeruginosa*) and negative droplets visually, since *E. coli* and *P. aeruginosa* droplets exhibited significantly different colony shapes. We classified droplets containing *P. aeruginosa* (positive) collected in the positive channel as true positives, while empty or *E*.*coli* ones (negative) that ended up in the positive channel were deemed false positives. Similarly, the negative droplets in the negative channel were counted as true negatives, whereas positive droplets in the negative channel were identified as false negative incidents. We calculated and employed the following metrics: i) **Accuracy** indicating the combined proportion of true positive and true negative sorting events relative to the total number of droplets. ii) **Sensitivity** which represents the true positive sorting events in relation to the total number of collected droplets with proteolytic colonies, which is calculated as the sum of true positives and false positives. iii) **Specificity** which measures the true negative events compared to the total number of empty droplets and droplets with *E. coli* (the sum of true negatives and false negatives). Throughout this optimization phase, we tested an emulsion composed of a mixture of *P. aeruginosa* droplets (1% of the total), *E. coli* droplets (10%) and empty remaining droplets. We covered a spectrum of droplet flow rates, corresponding to various droplet frequencies ranging from 20 to 150 droplets per second. Simultaneously, we explored sorting oil flow rates spanning from 0.6 to 3 mL/h. To maintain consistent spacing between droplets, we kept the spacing oils flow rates constant at 3 ·10^−2^ mL/h and 6 ·10^−2^ mL/h for the first and second spacing oil, respectively. As shown in **Figure 6**, our results reveal that the best levels of accuracy, selectivity, and sensitivity were accomplished when sorting oil flow rates were within the range of 1.8-3 mL/h and droplet frequencies of up to 50 Hz.

**Figure 6.**
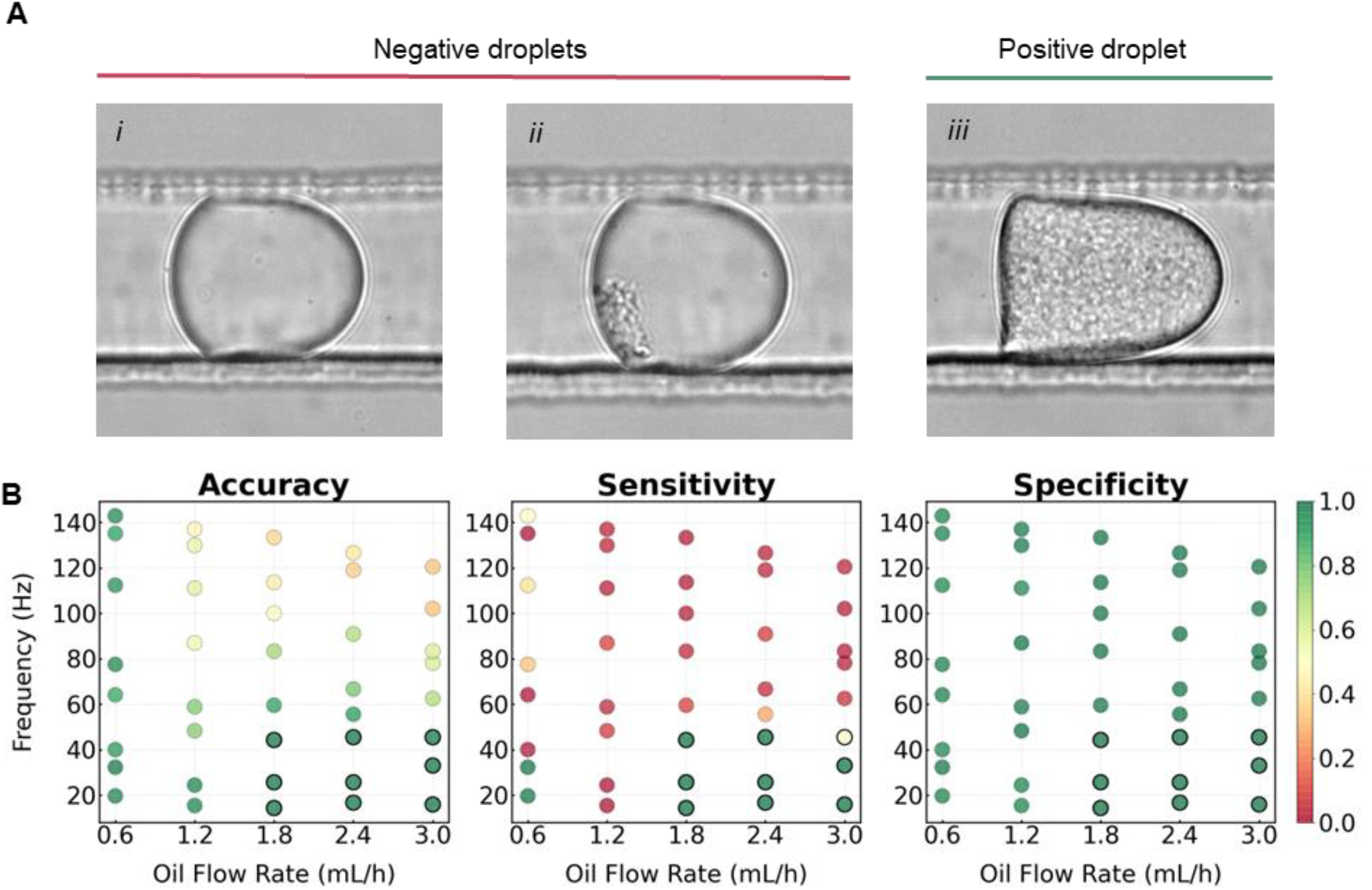
Impact of various flow rates of sorting oil on droplet sorting under microbial cultivation conditions. To mimic the application of the DPDS device towards the screening of an environmental sample, we tested a population of droplets comprising 2% *P. aeruginosa* droplets (positives) and 10% *E. coli* droplets (negatives). In Panel A, we reported the three types of droplets analyzed, i) empty droplet, ii) *E*.*coli* culture, iii) *P. aeruginosa* culture. We varied the throughput frequencies by adjusting the emulsion flow rate (6 ·10^−3^ - 4.8 ·10^−2^ mL/h), and in parallel we tested different sorting oil flow rates (0.6-3 mL/h). The three key parameters we examined are presented in Panel B: Sorting accuracy, sensitivity and specificity. The green dots marked with black contours highlight the flow rates of droplets and sorting oil to achieve 100% sorting efficacy.

**Figure 7.**
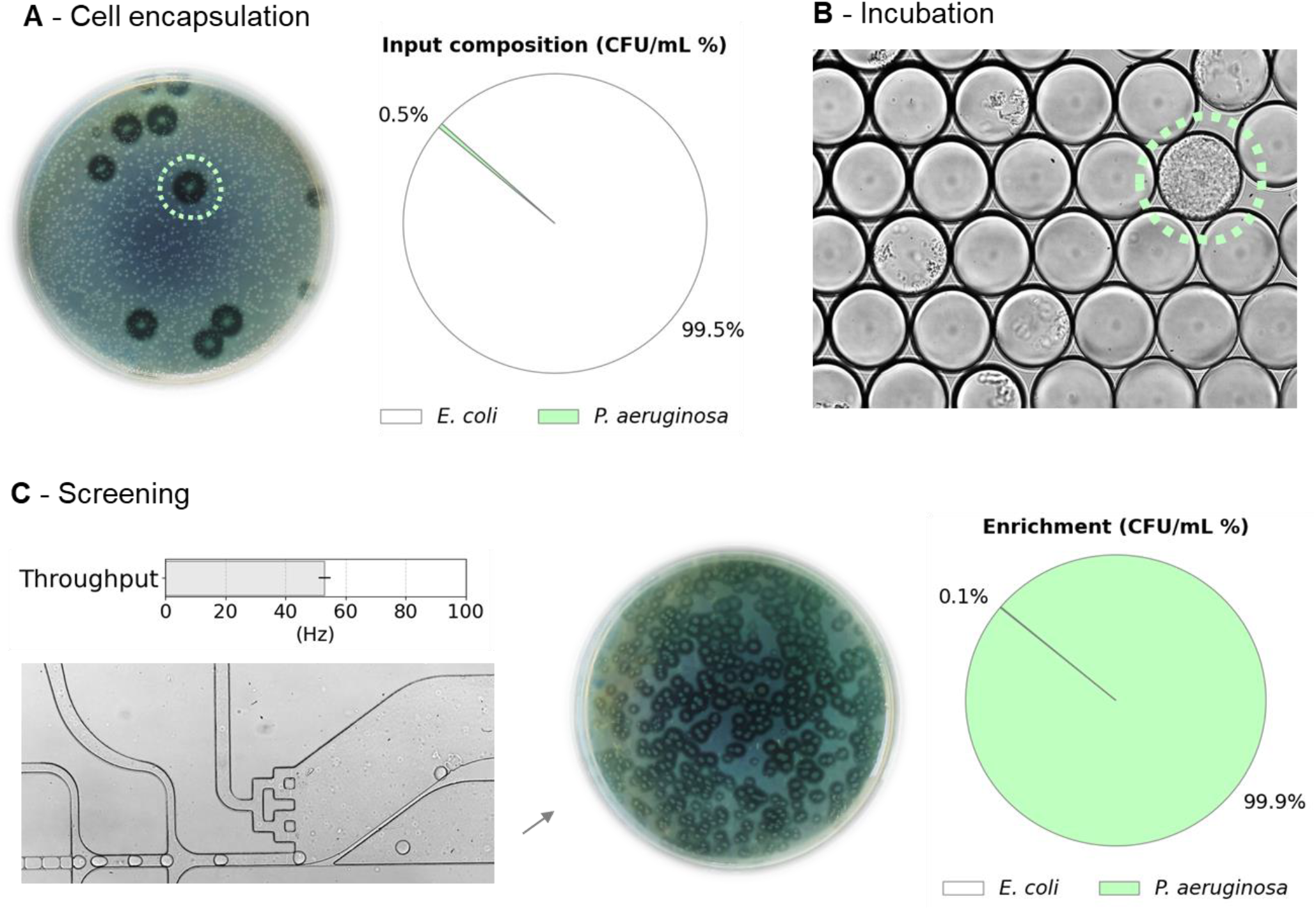
Enrichment of proteolytic strain from the mock 2-strain consortium. Panel A displays a Petri dish containing bacterial colonies present after the initial inoculum in the gelatin medium prior to the generation of droplets. It predominantly consists of *E. coli* colonies, with only a few colonies of *P. aeruginosa* indicated by a transparent halo. The relative abundance of *P. aeruginosa* in terms of CFU/ml was approximately 0.5% out of the total colonies enumerated. Droplets of 100 pL cultivation medium supplemented with 75 g/L gelatin were incubated at 40°C for 72 hours (Panel B) and then screened in the DPDS device as shown in Panel C. The Petri dish in the lower section represents the outlet collection of colonies predominately (>99%) formed by *P. aeruginosa*, distinguishable by the transparent halo around them.

### Enrichment of proteolytic strain *P. aeruginosa* from a mock microbial community emulating an environmental sample

To validate the accuracy of our novel passive sorter, we enriched positive droplets encapsulating single-cell originating colonies of the highly proteolytic *P. aeruginosa* strain from a synthetic microbial consortium. The emulsion primarily consisted of empty droplets (approximately 80%) to minimize the chance of having doublets or triplets of cells per droplet. Analysis of videos from sorting experiments provided insights into the accuracy of the enrichment, including the rate of false-negative and false-positive events. *P. aeruginosa* and *E. coli* were cultured separately overnight in flasks and then mixed to create a mock consortium. This mixture was suspended in an LB medium to achieve a concentration corresponding to approximately λ=0.2 and encapsulated in 100 pL microdroplets. After overnight cultivation, the emulsion was transferred to the DPDS device, and in 90 minutes, around ∼270 000 droplets were sorted and collected in sterile tubes. The enrichment experiments were conducted in triplicates. The aqueous phase of the collected and subsequently broken emulsion (positive droplets) was resuspended in sterile NaCl saline solution, then diluted 100 times before plating on agar medium supplemented with skimmed milk. Plates containing at least 500 colonies were chosen to count positive and negative colonies. The presence of active proteases, which can break down the milk substrate, was indicated by halos around positive bacterial colonies. Footage captured during the enrichment of this mock consortia is available in the Supporting Information SI section (**Video S2**).

The enrichment factors were determined using previously described methods by Baret et al.^29^ and Zinchenko et al.^30^. The Baret method (**a**) calculates enrichment (**η**) by comparing the final ratio of positive to negative colonies (ε1) after sorting with the initial ratio (ε0) before sorting. The Zinchenko method (**b**) calculates the enrichment factor (**η’**) by comparing the percentage of positive colonies to the total number (ε1’) with the initial percentage (ε0’).

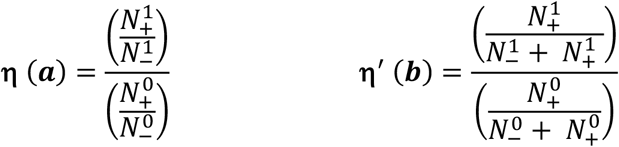

The values of factors we calculated were exceptionally high - approximately 4 ·10^5^ and 2 ·10^2^ - indicating that our approach effectively enriches proteolytic microorganisms.

**Table 1.**
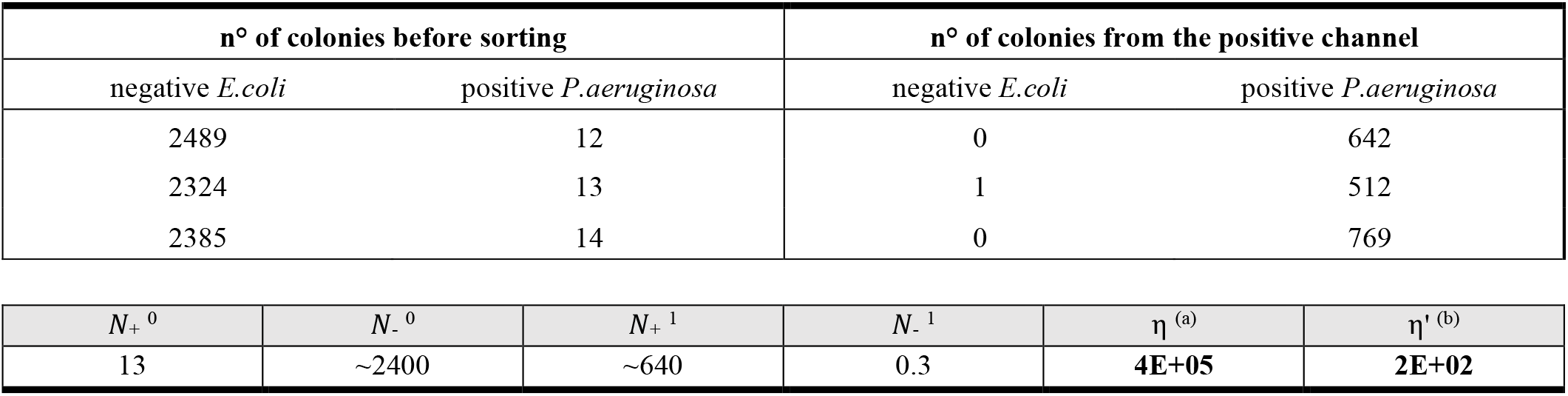
The enrichment values of the DPDS method. Bacterial suspensions were plated at a 100× dilution for the unsorted emulsion and for the emulsion after sorting.

## CONCLUSIONS

The application of accurate tools to screen microbial consortia holds great promise for the modern microbiology and biotechnology industry^31^. Today, the development of precise and fast screening of strains is still difficult, however the implementation of droplet microfluidics protocols can facilitate this process. Here, we described an approach based on single-cell encapsulation and subsequent droplet cultivation for screening microbial proteolytic activity. The enrichment of proteolytic microbes is performed in a novel chip device based on a decreased deformability of gelatin droplets due to proteolytic activity. Ultimately, we identified the optimal conditions that allowed us to passively sort up to 50 droplets per second. To enhance the efficiency of droplet sorting, we conducted a series of experiments to optimize several parameters. These included adjusting the volume of generated droplets to 100 pL, determining the optimum flow rates for droplets, spacing, and sorting oils, as well as identifying the gelatine concentration necessary for generating droplets while maintaining emulsion stability during incubation. Eventually, we achieved a consistent and high level of sorting accuracy, and next we focused on isolating droplets containing proteolytic microbes out of a mock consortium that mimics an environmental sample.

In contrast with the current Lab-On-a-Chip formats, the proposed enrichment platform does not require the use of expensive or not-commercially available fluorogenic substrates to detect protein degradation. The novel sorter design we introduced relies on a passive sorting mechanism based on the deformability properties of the cultivation medium used to generate picoliter droplets. Compared to other high-throughput methods used for screening proteases via fluorescence-based assays^20,32^, the simplicity of our method does not foresee the use of electronics or optomechanical equipment, making it more user-friendly. During passive sorting, an operator should focus exclusively on adjusting the combination of oil flows to execute precise sorting and high enrichment. Furthermore, when compared to other passive systems used for deformability-based screening of droplets, our method operates at significantly higher rates, as reported in Table 2. In contrast to the screening of agarolytic activity conducted by Muta et al.^24^, our device performs passive droplet sorting at a throughput two orders of magnitude higher. The double-rail layout proposed by Muta et al. works only if droplets flow at a relatively slow rate to effectively leap to the second rail, which directs them to the positive outlet. In general, rail chips might be less optimal for deformability-based sorting due to the inherent challenge of continuously compressing solid hydrogel droplets during the whole sorting operation, which might lead to clogging of the device, as well as droplet collisions and merging.

**Table 2.**
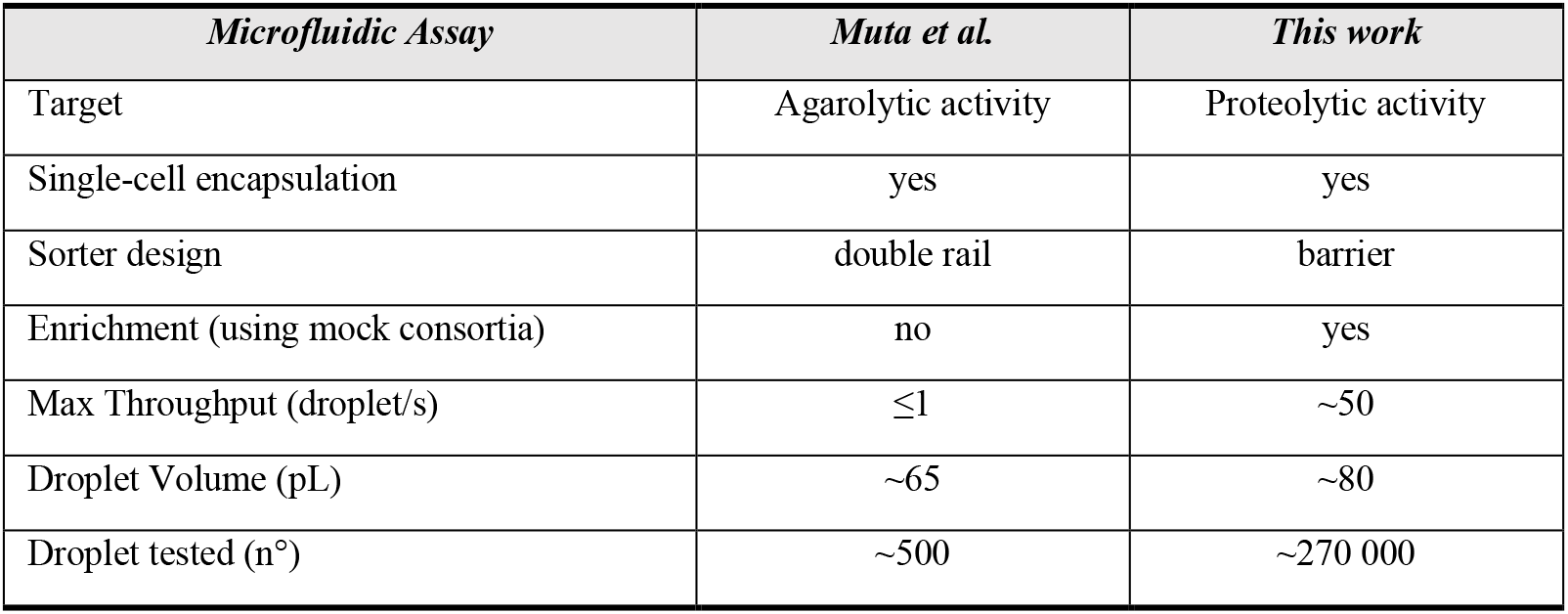
Comparison with other recently developed deformability-based sorting of droplets.

The gelatin-based strategy we propose is limited by the various temperatures needed during droplet generation and sorting. This requirement arises from the fact that the deformability of the gelatin medium changes with temperature. Future improvements can involve exploring alternative protease substrates, such as methacrylate gelatin, since this could potentially help address the temperature-related challenges.

Our droplet microfluidic protocol, designed for isolating proteolytic microbes, may have broader implications beyond its initial scope. The DPDS system can be used for screening hydrogels, to evaluate both their polymerization and degradation processes when exposed to microbial cells. This system is not only rapid but also cost-effective, making it an ideal solution for advancing other studies, such as the directed evolution of enzymes and detecting the secretion of proteases by single mammalian cells ^20^. Taken together, we believe that the DPDS system could significantly advance both research and practical applications within environmental microbiology. Notably, the DPDS system is ideally suited for monitoring proteolytic bacteria populations and improving our understanding and control of microbial interactions during, e.g., biofuel production and bioremediation processes. Additionally, the system aids in isolating rare microbial strains from diverse environments, essential for advancing industrial and environmental applications.

## ASSOCIATED CONTENT

### Supporting information

Detailed protocols of photolithographic microfabrication of molds of microfluidic devices, droplet chamber and incubation, and video captions.

- CAD file with designs of microfluidic chips (.dxf).
- An image of a profilometer scan of sorting module (Figure S1)
- Results of microbial growth optimization (Figure S4)
- Video S1 high viscosity gelatin droplets generation.
- Video S2-S3 passive sorting of 0% and 7.5% gelatin droplets.
- Video S4 of passive sorting during enrichment of 2-strain mix

## ACKNOWLEDGEMENTS

This research was funded by TEAM-NET programme of the Foundation for Polish Science, project no. POIR.04.04.00-00-14E6/18-00 as a part of Measure 4.4 of the 2014–2020 Smart Growth Operational Programme, EU and by National Science Centre, Poland (grant SONATA BIS no. 2023/50/E/ST4/00545). project. Research infrastructure used in the project was co-funded by the “Excellence Initiative – Research University (2020-2026)” IDUB programme via Action I.4.2 “Fund for the Renovation and Development of Research Infrastructure.” IDUB small grants funded by the University of Warsaw also contributed to the execution of this research work (grant no. 01/14-01-00/2023). We would like to thank Marta Zapotoczna for the access to the plate reader, Julia Karbowska for the useful advice in the initial stage of the project and Lukasz Dziewit and Mikolaj Wolacewicz for help with 16S sequencing the proteolytic reference strain used in this study. Figures of EPDS workflow and scheme of dynamic droplet incubation in the SI were prepared with BioRender.com.

