## Supporting information for "Passive droplet microfluidic platform for high-throughput screening of microbial proteolytic activity"

**Fabrication of microfluidic devices.** The microfluidic devices used in this work were designed in AutoCAD (Autodesk), and the master molds were fabricated by standard photolithography and soft lithography[1]. To fabricate PDMS devices, the following steps were executed: first, master molds containing designed structures were made via photolithography, and then the replicas containing the channels were produced by pouring PDMS onto the master molds.

**Photolithography.** Microfluidic molds were created on 3-inch silicon wafers (Microchemicals) using high-resolution acetate masks (Microlithography Services) and SU-8 series photoresist patterning (Kayaku Advanced Materials). The SU-8 spin-coated wafers were then exposed to the UV using an MJB4 mask aligner (SÜSS MicroTec)[2]. The CAD designs of the droplet generation device and DPDS (Deformability-based Passive Droplet Sorter) sorter module were attached as a supplementary .dxf file. The thickness of the structures (corresponding to the depth of channels in the final microfluidic devices) was measured using a GT-Contour profilometer (Bruker) and 5x objective. The profile of the droplet sorting module is presented in Figure S1.

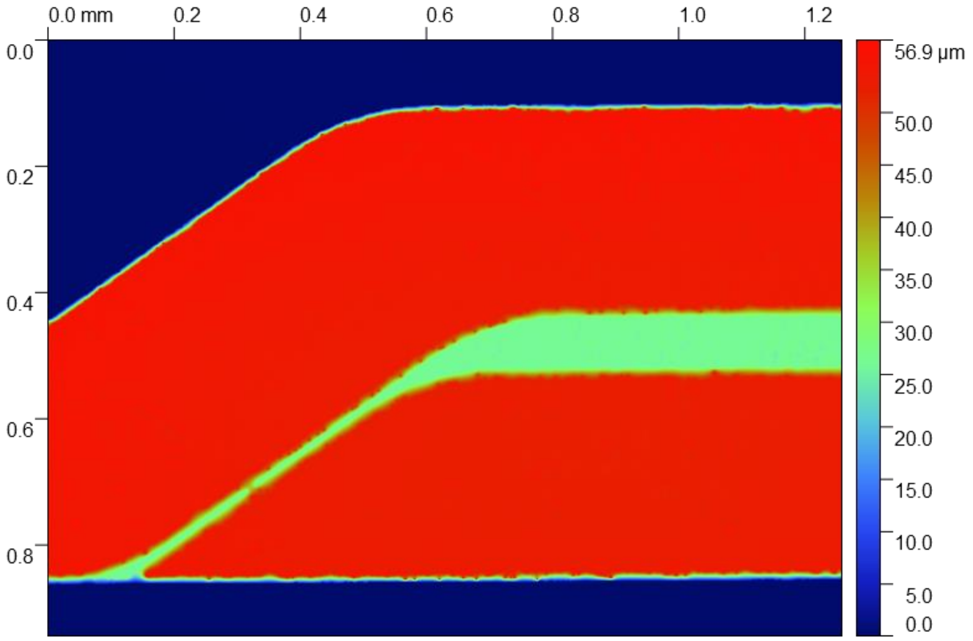


**Figure S1. Profilometry scan.** The picture highlights the barrier and showcases the double-layered layout of the DPDS sorter.

**Table S1. Fabrication protocol.** Settings for the photolithography of the DPDS sorter module.

| | **Fabrication step (no. of a layer)** | |
|---|---|---|
| | **1st layer** | **2nd layer** |
| Nominal thickness | 30 µm | 30 µm (60 µm final thickness) |
| Resist used | SU-8 2025 | SU-8 2025 |
| Spin coating speed | 1st step: 10 sec, 500 rpm<br>2nd step: 30 sec, 3000 rpm | 1st step: 10 sec, 500 rpm<br>2nd step: 30 sec, 4500 rpm |
| Pre-baking | 1 min at 65ºC<br>5 min at 95ºC | 1 min at 65ºC<br>5 min at 95ºC |
| Exposure (at ~12.8 mW $cm^{-2}$) | 12 sec | 12 sec |
| Post-baking | 1 min at 65ºC<br>5 min at 95ºC | 1 min at 65ºC<br>5 min at 95ºC |
| Development in the beaker filled with 30-50mL of PGMEA (propylene glycol methyl ether acetate, Sigma-Aldrich) | n.a. | Approximately 5-10 minutes until all uncured SU-8 was removed from the wafer |
| Measured range of thicknesses | 28 µm | 30 µm |

**Soft lithography.** To generate a single microfluidic device, approximately 30 grams of PDMS (Sylgard 184, Dow) are weighed in a plastic cup. The curing agent was added and mixed at a 10:1 (w/w) ratio, and the liquid PDMS was then degassed in a vacuum chamber. PDMS is poured on the SU-8 master wafer, placed in Petri dish, and cured in the oven at 70°C for 4 hours. Once the PDMS is solidified, the device is peeled off from the mold, and the holes for inlets and outlets are punched using a 1 mm biopsy puncher (Kai Medical). The flow-focusing droplet generators and the DPDS sorter chips were bound to glass slides (VWR) using an automated plasma system (Zepto, Diener Electronics). The plasma treatment was conducted as follows: first, a vacuum of 0.35 mbar was generated. Then, oxygen was introduced into the chamber for one minute at a pressure of 0.5 mbar. Finally, the plasma process was initiated at 30% power for 45 seconds.
Hydrophobic modification of the chips was carried out by flushing the devices with a 1% solution of trichloro (1H,1H,2H,2H perfluorooctyl)-silane (Sigma-Aldrich) in Novec HFE-7500 oil (3M) and baked on a hot plate at 80°C for 30 minutes to evaporate fluorocarbon liquid.

**Fabrication of storage chambers for droplet incubation.** The chambers were made according to the protocol published by Neun et al.[3] A biopsy punch (Kai Medical) was used to make 1-mm holes on the bottom tip and the side of a 0.5 ml Eppendorf tube. The tube lid was then glued to a 1-mm thick glass slide with cyanoacrylate glue (PR 1500, 3M). Two pieces of 20-cm-long Teflon (PFTE) tubing (0.4mm I.D., 0.9mm O.D, Bola Bohlender) were then inserted into the holes and glued to the surface of an Eppendorf tube. During the droplet generation, the flow-focusing chip was connected to the upper tubing, allowing droplets to be collected in the upper section of the tube - due to their natural buoyancy compared to perfluorinated oil. After incubation, to sort the droplets, the direction of flow was reversed, and the oil was pumped through the side tubing, and moving the droplets towards the DPDS sorter module via the top tubing.

**Droplet oxygenation during incubation in storage chambers.** Emulsions were incubated for 72 hours by dynamic droplet incubation (DDI), which enhances microbial growth and activity[4]. The setup used by us in this study is presented in Fig. S2. Measures were taken to remove any bubbles present in the tubing. The schematic representation of the setup employed for oxygenating the droplets in the experimental workflow. It highlights the crucial role of the peristaltic pump and bubble trap that simultaneously allowed for oxygen delivery and prevented air bubble formation in the incubation chamber. The pump operates using Tygon tubing of 0.38 mm I.D. and 0.90 mm O.D. (Ismatec), such pieces of tubing were connected and glued with PTFE tubing (0.5 mm I.D. and 1 mm O.D., Bola Bohlender) of the incubation chambers. The layout of this sealed system (Figure S2) was completed by putting two sides of the tubing into a so-called bubble trap, a 1.5 mL Eppendorf tube containing 5% RAN fluorosurfactant in HFE-7500.

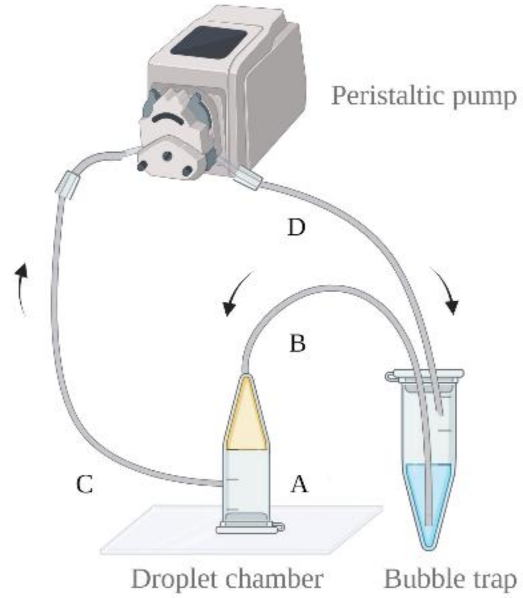


**Figure S2. Oxygenation system layout.** The first 1.5 ml Eppendorf tube was glued to a glass slide and used as a droplet chamber (**A**). Then one of the tubing was glued to the top part of the chamber (**B**). A second tubing was inserted closer to the lid (**C**), and both fragments of tubing were securely connected via a piece of PTFE tubing to a peristaltic pump (Reglo ICC, Ismatec) using adhesive glue. The other end of the tubing coming from the pump was glued on the lid of a second Eppendorf tube filled with 5% RAN 008-FluoroSurfactant solution in filtered Novec HFE-7500 (**D**). This arrangement ensured a continuous flow of the oil phase in a closed system. Oil flow of oil is directed from the top of the tube, where droplets accumulate, to the lower inlet of the chamber at 0.6 ml/h. As a result, the emulsion was held in place by buoyancy, and droplets did not leak out of the chamber into the oil solution.

**Microbial isolation of reference strain.** The proteolytic reference strain utilized in this study was isolated from the sludge of a wastewater treatment plant located in Wołomin, Mazovia region, Poland. The sludge underwent resuspension in a physiological solution within 0.1 L flasks and was then subjected to shaking for 1 hour at 200 rpm at 30°C. Subsequently, the sample was diluted and spread onto skimmed milk agar medium (30g/L), known for its selectivity towards proteolytic activity. Further identification through 16S sequencing confirmed that our strain was identified as *Pseudomonas aeruginosa*.

**Microbial growth optimization.** We optimized microbial growth inside droplets to analyze the proteolytic activity of microcultures originating from individual bacteria as soon as possible to avoid any possible issues, e.g., merging of droplets or osmosis leading to negative droplet shrinkage. To achieve rapid microbial growth, we analyzed the growth curve of the proteolytic reference strain in various cultivation media, different flow rates of incubation oil, and additional oxygenation of the oil within the storage chambers (bubbling). In Figure **S4**, we have presented the impact of microbial growth in different media on two gelatin concentrations.

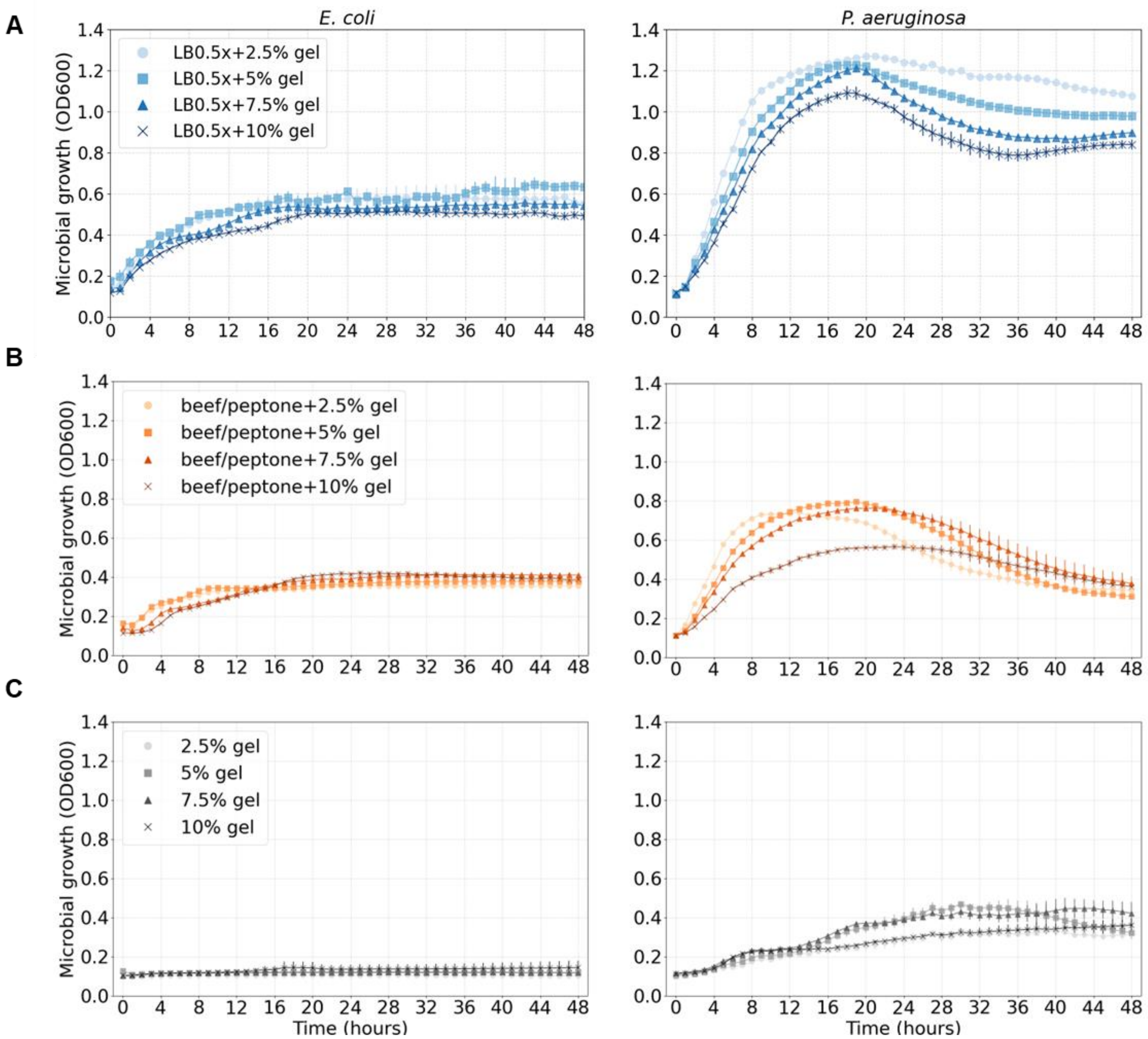


**Figure S4. Cultivation medium optimization and influence of gelatin concentration over microbial growth.** We first measured the microbial growth of the reference strains we used in different cultivation media supplemented with different concentrations of gelatine. In Panel **A,** gelatin was resuspended in a diluted form of Luria-Bertani medium (LB0.5x). In Panel **B,** we tested the recommended medium used for screening of proteolytic activity composed of peptone (5 g/L) and beef extract (3 g/L). Finally, in Panel **C,** we measured growth in pure gelatine solutions. Microbial growth of proteolytic reference strain was analyzed by an automatic plate reader (Synergy HTX, BioTek) every hour for 48 hours, at minimal orbital shaking speed and 40°C temperature.

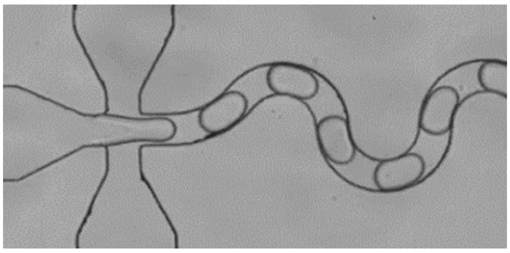

**Video S1. Generation of gelatin droplets.** Emulsions were generated in a 50 x 50 µm flow-focusing device. The maximum throughput we achieved by using the final concentration of gelatine of 75 g/L to generate droplets (100 pL) was around 1.6 kHz. This video was recorded at 10,000 frames per second (FPS) and 1/20,000 shutter speed.

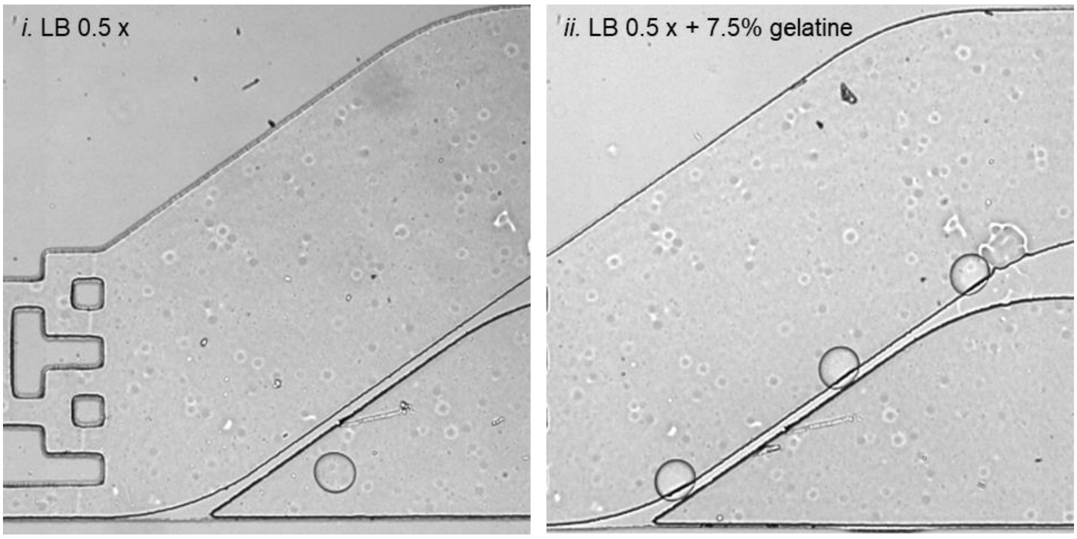


**Video S2. Passive sorting of droplets with different gelatin concentrations.** The passive sorting of droplets with different concentrations was tested, and emulsions were separately screened in the DPDS device. Videos were recorded at 5,000 FPS and 1/50,000 shutter speed. The sorting frequency was approximately 50 Hz.

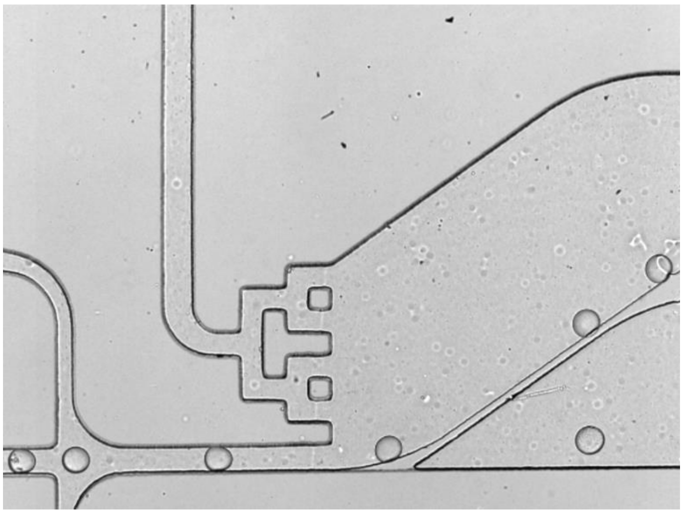

**Video S3. Passive sorting during enrichment of 2-strain mix.** The sorting process was applied to an emulsion composed of the mock microbial community of *E. coli* and *P. aeruginosa*. Oil flow was directed from left to right. The module with barrier sorted the droplets with *P. aeruginosa* into the positive channel (lower section). All the other droplets, empty and *E. coli* ones, were derailed to the negative channel (upper section). This video was recorded at 5,000 FPS and 1/50,000 shutter speed. The sorting frequency was approximately 50 Hz.
